# Indirect Correlative Light and Electron Microscopy (*iCLEM*): A Novel Pipeline for Multiscale Quantification of Structure from Molecules to Organs

**DOI:** 10.1101/2023.05.15.540853

**Authors:** Heather L. Struckman, Nicolae Moise, Bieke Vanslembrouck, Nathan Rothacker, Zhenhui Chen, Jolanda van Hengel, Seth H. Weinberg, Rengasayee Veeraraghavan

## Abstract

Correlative light and electron microscopy (*CLEM*) methods are powerful methods which combine molecular organization (from light microscopy) with ultrastructure (from electron microscopy). However, CLEM methods pose high cost/difficulty barriers to entry and have very low experimental throughput. Therefore, we have developed an *indirect* correlative light and electron microscopy (*iCLEM*) pipeline to sidestep the rate limiting steps of CLEM (i.e., preparing and imaging the same samples on multiple microscopes) and correlate multiscale structural data gleaned from separate samples imaged using different modalities by exploiting biological structures identifiable by both light and electron microscopy as intrinsic fiducials.

We demonstrate here an application of *iCLEM*, where we utilized gap junctions and mechanical junctions between muscle cells in the heart as intrinsic fiducials to correlate ultrastructural measurements from transmission electron microscopy (TEM), and focused ion beam scanning electron microscopy (FIB-SEM) with molecular organization from confocal microscopy and single molecule localization microscopy (SMLM). We further demonstrate how *iCLEM* can be integrated with computational modeling to discover structure-function relationships. Thus, we present *iCLEM* as a novel approach that complements existing CLEM methods and provides a generalizable framework that can be applied to any set of imaging modalities, provided suitable intrinsic fiducials can be identified.

## Introduction

“*Form follows function*”, as architect Louis Sullivan famously stated. Thus, biologists have long studied structure in order to understand how biological systems function. In terms of microscopy, this presents a multiscale imaging challenge ranging from the molecular level up to the macroscopic scales of organs and organisms. Historically, electron microscopy has offered exquisite spatial resolution and the ability to visualize the ultrastructure of cells and tissues, whereas light microscopy has enabled scientists to visualize specific biomolecules *in situ*, albeit with little to no structural context and limited spatial resolution. Thus, correlative light and electron microscopy (CLEM) methods, which enable specific biomolecules to be visualized within their ultrastructural context, stand to revolutionize the life sciences. However, CLEM methods entail exorbitant expenses, and pose major technical challenges due to the disparate sample chemistries demanded by light and electron microscopy modalities and the challenges associated with precisely registering images of nanoscale regions viewed through separate microscopes. These factors have severely limited the usability of CLEM for many researchers as well as its multiscale investigative capabilities and experimental throughput, thereby precluding robust assessment of intra- and inter-sample heterogeneities necessary to capture subtle and/or rare phenomena. Thus, new approaches are needed to complement the strengths of CLEM and address use cases, where greater throughput is needed.

To this end, we have developed an *indirect* correlative light and electron microscopy (*iCLEM*) pipeline to sidestep the rate limiting steps of CLEM (i.e., preparing and imaging the same samples on multiple microscopes) and instead, correlate multiscale structural data gleaned from separate samples imaged using different modalities by utilizing structures within biological samples that can be visualized by both light and electron microscopy as intrinsic fiducials. We demonstrate here, using cardiac muscle tissue as a test case, how gap junctions and mechanical junctions between muscle cells, which are readily visualized via both light and electron microscopy, can be exploited as intrinsic fiducials to correlate ultrastructural measurements from transmission electron microscopy (TEM), and focused ion beam scanning electron microscopy (FIB-SEM) with measurements of molecular organization gleaned from confocal microscopy and single molecule localization microscopy (SMLM; specifically stochastic optical reconstruction microscopy [STORM]). By giving up direct image-to-image correlation and embracing collection of large datasets (hundreds to thousands of images per sample group per technique), *iCLEM* captures multiscale structural properties and heterogeneities. We also demonstrate how this experimental pipeline can be readily integrated with computational modeling to uncover the functional implications of observed structural properties. Thus, *iCLEM* complements existing CLEM methods and provides a generalizable framework that can be applied to any set of imaging modalities, provided suitable intrinsic fiducials can be identified.

## Materials and Methods

All animal procedures were approved by The Ohio State University Institutional Animal Care and Use Committee and performed in accordance with the Guide for the Care and Use of Laboratory Animals published by the U.S. National Institutes of Health (NIH Publication No. 85-23, revised 2011).

### Sample Collection

#### Whole Heart Isolation

Adult wild-type (WT) mice (C57BL/6 background), aged 12-34 weeks, purchased from Jackson Laboratories (Cat. #000664), were used for tissue studies. They were anesthetized with 5% isoflurane + 100% oxygen (1L/min) and once unconsciousness occurred, anesthesia was maintained within 3-5% isoflurane + 100% oxygen (1L/min). After the animal was in a surgical plane of anesthesia, the heart was excised and immediately submerged in cold electrolyte solution (containing in mM: NaCl 140, KCl 5.4, MgCl_2_ 0.5, dextrose 5.6, HEPES 10; pH adjusted to 7.4) to remove any remnants of blood prior to cryopreservation (for immunofluorescence), fixation (for electron microscopy), or Langendorff perfusion (for myocyte isolation) as previously described(Mezache, et al., 2020; Moise, et al., 2021; Struckman, et al., 2020).

#### Preservation of Whole Hearts

Hearts were cryopreserved in optimal cutting temperature (OCT) compound for cryosectioning and use in immunofluorescence studies as previously described(Mezache, et al., 2020; Moise, et al., 2021; Struckman, et al., 2020). Briefly, hearts were dried to remove clinging water droplets prior to cryopreservation. A base layer of OCT compound (Fisher Health Care, Houston, TX, USA) was cooled in the cryomold with dry ice until partially solidified before placing the heart (to secure its position), then the remainder of the mold was filled with OCT compound. The cryomold was floated on liquid nitrogen to ensure uniform freezing using vapor phase nitrogen and once frozen, stored at -80 °C over the long term. OCT blocks were transferred to -20°C for 12-24 hours prior to cryosectioning. Immunofluorescence studies were performed on 5 µm sections fixed in 2% paraformaldehyde (PFA; Millipore Sigma, St. Louisa, MO) for 5 minutes at room temperature (RT).

For transmission electron microscopy (TEM) studies in cardiac tissue, ∼0.5mm cubes of tissue were fixed overnight in 2.5% glutaraldehyde (Electron Microscopy Sciences, Hatfield, PA USA) at 4°C followed by a 5 min wash in PBS and storage in PBS at 4°C for further processing. Volume electron microscopy data used here were derived from previously published work(Vanslembrouck, et al., 2018). Briefly, wild-type mice (aged 30–43 weeks) were anesthetized by intraperitoneal injection of ketamine/xylazine (70 mg of ketamine and 10 mg of xylazine per kg of body weight) and perfused for 2 min with PBS containing heparin (20 units/ml), followed by 10 mins of perfusion with 2% paraformaldehyde (PFA) + 2.5% glutaraldehyde in 0.15 M cacodylate buffer, pH 7.4, for 10 min. Next, heart muscle tissue was isolated and fixed overnight by immersion in 2% PFA + 2.5% glutaraldehyde in 0.15 M cacodylate buffer, pH 7.4. Finally, samples were thoroughly washed in 0.15 M cacodylate buffer, pH 7.4, before small blocks from regions of interest were dissected out to proceed with the staining protocol.

#### Preservation of Isolated Cardiac Muscle Cells

To isolate cardiac muscle cells for light microscopy studies, Langendorff-perfused hearts were enzymatically digested using Liberase™ TH (Roche, Basel, Switzerland) enzyme in Ca^2+^-free electrolyte solution. The upper (atria) and lower chambers (ventricles) were dissected after digestion and separately minced in electrolyte solution containing 2% Bovine Serum Albumin (BSA; Sigma-Aldrich, Bornem, Belgium), dispersed by gentle agitation, and filtered through a nylon mesh. Cardiac muscle cells were then resuspended in low Ca^2+^ electrolyte solution (containing 0.1 mM CaCl_2_). Cells were plated on laminin-coated glass coverslips, fixed for 5 min at RT with 2% paraformaldehyde (PFA), washed in phosphate-buffered saline (PBS) (3 × 10 min at RT) and stored in PBS at 4°C until ready for immunolabeling.

### Sample Preparation

#### Immunolabeling for Light Microscopy

Cryosections (5 μm thickness) of cardiac tissue were collected on glass slides (for confocal microscopy) or acid-washed, poly-lysine coated #1.5 glass coverslips (Electron Microscopy Sciences, Hatfield, PA; for single molecule localization microscopy) and fixed with 2% PFA in PBS (5 min at RT) followed by washes with PBS (3 x 10 min at RT). Isolated cardiac muscle cells adhered to #1.5 glass coverslips using laminin were similarly fixed in PFA and washed in PBS. Samples (either fixed tissues or cells) were permeabilized with 0.2% Triton X-100 in PBS (15 min at RT) and incubated with blocking agent (1% bovine serum albumin, 0.1% triton in PBS for 2 hrs at RT). Samples for confocal microscopy were labeled with primary antibodies (overnight at 4°C), washed in PBS (3 x 5 min at RT), labeled with secondary antibodies (2 hrs at RT) and washed again in PBS (3 x 5 min at RT) before being mounted in Prolong Gold (Invitrogen by Thermo Fisher Scientific, Grand Island, NY, USA) and allowed to cure for 48 hours at RT before imaging. Samples for STochastic Optical Reconstruction Microscopy (STORM) were processed similarly with the following minor changes. Washes before and after secondary antibodies were performed using the blocking agent diluted 10x in PBS (final concentration 0.1% bovine serum albumin, 0.01% triton in PBS) in place of PBS. Rather than being mounted on glass slides, labeled samples on 20 mm coverslips were post-fixed in 2% PFA (5 minutes at RT) followed by PBS washes (3 x 5 min at RT), then optically cleared in Scale U2 buffer (4M urea + 30% glycerol + 0.1% triton in water at 4°C) for 48 hours before imaging.

For all immunolabeling studies, well-validated custom and commercial primary antibodies against proteins of interest were carefully selected and experimental parameters rigorously optimized to ensure high fidelity labeling: Connexin 43 (Cx43; mouse monoclonal antibody, Cat. # MAB3067, EMD Millipore Corp., Darmstadt, Germany; rabbit polyclonal antibody, Cat. # C6219, Millipore Sigma, St. Louis, MO), N-cadherin (N-cad; mouse monoclonal antibody, Cat. # 610921, BD Biosciences, San Jose, CA), desmoplakin (Dsp; rabbit polyclonal antibody, Cat. # ab71690, Abcam, Waltham, MA), inward-rectifier potassium channel (K_ir_2.1; rabbit polyclonal antibody, Cat. # APC-159, Alomone labs, Jerusalem, Israel), cardiac voltage-gated sodium channel (Na_V_1.5; custom rabbit polyclonal antibody(Veeraraghavan, et al., 2018), sodium/potassium ATPase (NKA; a validated monoclonal custom antibody(Chen, et al., 2022), ryanodine receptor (RyR2; mouse monoclonal antibody, Cat. # MA3-916, ThermoFisher, Grand Island, NY, USA), connexin 40 (Cx40; rabbit polyclonal antibody; Cat. # 36-4900, ThermoFisher Scientific, Waltham, Massachusetts, USA), sodium channel β subunit (β1; custom rabbit polyclonal antibody(Veeraraghavan, et al., 2018), and nuclear stain (Hoechest 33342 nucleic acid stain, Cat. # H3570, ThermoFisher Scientific, Waltham, Massachusetts, USA). For confocal microscopy, samples were labeled with goat anti-mouse and goat anti-rabbit secondary antibodies conjugated to Alexa 488 and Alexa 568 (1:4000; ThermoFisher Scientific, Grand Island, NY). For single molecule localization microscopy, samples were labeled with goat anti-mouse Alexa 647 (1:1000) and goat anti-rabbit Biotium CF 568 (1:2000) secondary antibodies (ThermoFisher Scientific, Grand Island, NY, USA). Where needed, simultaneous labeling with two primary antibodies from the same animal background was accomplished by completing the labeling protocol with one set of primary and secondary antibodies followed by blocking with a donor secondary antibody (not conjugated to a fluorophore) to block any remaining unbound primary antibodies. A second round of labeling was then performed with a new combination of primary and secondary antibodies.

#### Staining for Electron Microscopy

Once preserved as described above, TEM samples were post-fixed with 1% osmium tetroxide (Ted Pella Inc., Redding, CA, USA) and then *en bloc* stained with 1% aqueous uranyl acetate (Ted Pella Inc., Redding, CA, USA), dehydrated in a graded series of ethanol, and embedded in Eponate 12 epoxy resin (Ted Pella Inc., Redding, CA, USA). Ultrathin sections (70 nm) were cut with a Leica EM UC7 ultramicrotome (Leica microsystems Inc., Deerfield, IL), collected on copper grids, and finally, stained with Reynold’s lead citrate and 2% uranyl acetate. Samples for volume electron microscopy were prepared as previously described(Vanslembrouck, et al., 2018). Tissue-blocks were post-fixed by incubation in 1% osmium (Electron Microscopy Sciences, Hatfield, PA) + 1.5% potassium ferrocyanide (Electron Microscopy Sciences, Hatfield, PA) in 0.15 M cacodylate buffer, pH 7.4. Once fixed, tissue blocks were washed in double-distilled H2O (ddH2O; Merck, Darmstadt, Germany), and stained with 1% heavy weight tannic acid (HWTA; Electron Microscopy Sciences, Hatfield, PA) in ddH2O in four consecutive steps, refreshing the HWTA after each step. Next, samples were incubated in 1% osmium in ddH2O followed by washing in ddH2O. This was followed by incubation in 1% uranyl acetate (UA; EMS) and another wash with ddH2O. After the final washes, samples were dehydrated using solutions of 50, 70, 90, 2×100% ethanol and next placed in 100% acetone. Embedding in Spurr’s resin (Electron Microscopy Sciences, Hatfield, PA) was performed by incubating the tissue blocks first in 50% Spurr’s in acetone and next in 100% Spurr’s, refreshing the solution at least three times. After this, samples were placed in Beem capsules containing fresh resin and polymerized overnight at 60 °C. All staining, dehydration and embedding steps, except for the Walton’s Lead were performed using a Pelco Biowave Pro Microwave Tissue Processor (TedPella Inc, Redding, CA USA).

### Image Collection

#### Confocal Microscopy

Confocal imaging was performed using an A1R-HD laser-scanning confocal microscope equipped with four solid-state lasers (405, 488, 560, and 640 nm, 30 mW each), a 10x/0.45 numerical aperture (NA) air objective, a 63x/1.4NA oil immersion objective, two GaAsP detectors, and two high sensitivity photomultiplier tube detectors (Nikon, Melville, NY, USA). Images were collected as single planes (for survey views only) or z-stacks (for all data used for quantitative analyses) with sequential spectral imaging (line-wise) in order to avoid spectral mixing. In addition, images were collected with Nyquist sampling (or greater) and post-processed with 3D deconvolution, as previously described(Mezache, et al., 2020; Moise, et al., 2021; Struckman, et al., 2020). Resolution achieved was in the 200 – 300 nm range.

#### Single Molecule Localization Microscopy

STochastic Optical Reconstruction Microscopy (STORM) was performed using a Vutara 352 microscope (Bruker Inc, Middleton, WI, USA) equipped with widefield illumination, a 60x magnification/1.3NA water immersion objective, multi-focus imaging (biplane) for axial position detection, and fast scientific complementary metal–oxide–semiconductor (sCMOS) camera achieving 20 nm lateral and <50 nm axial resolution. Individual fluorophore molecules were localized with a precision of 10 nm. Registration of the two color channels was accomplished using localized positions of several TetraSpeck Fluorescent Microspheres (ThermoFisher Scientific, Carlsbad, CA, USA) scattered throughout the field of view. Three dimensional images of *en face* IDs were collected at 150 frames/z-step.

#### Transmission Electron Microscopy (TEM)

TEM was performed on a FEI Tecnai G2 Spirit transmission electron microscope (Thermo Fisher Scientific, Waltham, MA) operating at 80keV, and a Macrofire (Optronics, Inc., Chelmsford, MA) digital camera and AMT image capture software.

#### Volume Electron Microscopy

Resin-embedded samples were trimmed and mounted on aluminum specimen pins (Gatan, Pleasonton, CA USA) using conductive epoxy (Circuit Works, Santa Monica, CA USA). Next, specimens were trimmed to a pyramid shape using an ultramicrotome and coated with an 8 nm platinum layer (Quorum Q150T ES sputter coater, Quorumtech, Laughton, UK). Focused ion beam scanning electron microscopy (FIB-SEM) imaging was performed at the VIB Bioimaging Core (Ghent University, Ghent, Belgium) using a Zeiss Crossbeam 540 with Atlas5 software. First, the SEM was used at 15 kV to localize an area containing the structure of interest, in this case, a specialized region of electromechanical contact between adjacent heart muscle cells known as an intercalated disc (ID). Subsequently, the FIB-SEM was set up to section at 5 nm and image at 1.5 kV using the Zeiss ESB detector. Registration of the datasets was performed using IMOD software (http://bio3d.colorado. edu/imod/)(Vanslembrouck, et al., 2018).

### Image Analysis

#### Spatial Analysis of Digital Images

Confocal images were analyzed using our previously published point process-based spatial analysis pipeline, Spatial Pattern Analysis using Closest Events (SPACE)(Bogdanov, et al., 2021). This pipeline, implemented in Matlab (Mathworks Inc, Natick, MA) uses distance transformations to efficiently measure the nearest neighbor distances for segmented immunosignals for a given biomolecules in relation to sub-cellular landmarks (eg. nuclei, cell periphery) or a second, co-labeled biomolecule. By comparing the distribution of observed nearest neighbor distances with that predicted under complete spatial randomness, quantitative measurements are obtained for the strength of non-random attraction / repulsion between co-labeled molecules / structures. Statistical comparisons between pairs of nearest neighbor distance distributions were accomplished using the 2 sample Kolmogorov-Smirnov test.

#### Analysis of Single Molecule Localization Data

Given that our research questions focused on the spatial relationships between clusters of biomolecules rather than individual molecules, we identified clusters of localizations and applied spatial statistical analysis at the cluster level. To this end, single molecule localizations were analyzed using our previously published STORM-based Relative Localization Analysis (STORM-RLA(Veeraraghavan & Gourdie, 2016)), a machine learning-based cluster analysis approach that uses density-based spatial clustering for applications with noise (DBSCAN) as the underlying clustering algorithm. By fitting surfaces to detected clusters, STORM-RLA calculates nearest neighbor distances between clusters of co-labeled biomolecules. Distributions of these nearest-neighbor distances can then be subjected to spatial statistical analysis as detailed above.

#### Morphometric Analysis of Electron Micrographs

Morphometric quantification was performed manually using ImageJ (National Institutes of Health, http://rsbwe b.nih.gov/ij/) as previously described(Struckman, et al., 2023). Measurements were performed on sets of TEM images of the same structures collected at different magnifications to capture properties at different spatial scales. Morphometric assessments were obtained from volume electron microscopy data by manually segmenting images using Microscopy Image Browser (MIB; http://mib.helsinki.fi/) to annotate structures of interest and computationally reconstructing them using Imaris (http://www.bitplane.com/imaris/imaris).

### Integration of Structural Measurements into Computational Models

In order to probe the functional implications of structural measurements gleaned from our light and electron microscopy studies, we integrated them into finite element models of the structures of interest, in our case, the cardiac intercalated discs. Following our previously described approach for mesh generation(Moise, et al., 2021), distributions of ultrastructural measurements derived from electron microscopy were sampled to construct populations of finite element models of intercalated discs. Notably, these meshes contained representations of gap junctions and mechanical junctions, which served as our intrinsic fiducials. Using our recently described approach(Struckman, et al., 2023), distributions of target proteins relative to these fiducials, which were derived from light microscopy studies, were sampled to populate the meshes with target proteins. The complete finite element models of intercalated discs, thus generated, were then integrated into functional models of electrical signal propagation through the heart to uncover structure-function relationships.

## Results

### The *Indirect* Correlative Light & Electron Microscopy (*iCLEM*) Workflow

Implementing *iCLEM* requires careful advance planning of all phases of the workflow including sample collection/handling, image collection, image analysis, and data correlation (Figure 1). The first step in this process is to plan the multiscale experimental / analytical framework by defining the specific measurements needed to address the research question(s) at hand and working backwards to select in tandem both the imaging modalities to be used and the intrinsic landmarks to be exploited to correlate measurements across imaging modalities (Figure 1A). It should be noted here that imaging modalities are selected based on their ability to resolve structures of interest and discriminate them from other structures (background) as well as the ability to capture sufficiently large regions of the sample’s volume to collect the necessary amount of observations. With these selections made, the subsequent steps of sample collection and preparation and image collection (Figure 1B-D) are planned as typically done for imaging studies using a single imaging modality with specific additions / exceptions are noted below. Next, image analysis approaches are selected to extract the necessary information from image data, which can best exploit / accommodate the unique properties of specific imaging modalities employed (Figure 1E). Last but not least, a plan is developed for correlating measurements across spatial scales and different imaging modalities by exploiting the structural fiducials for both correlation and cross-validation (Figure 1F, Figure 2).

**Figure 1.**
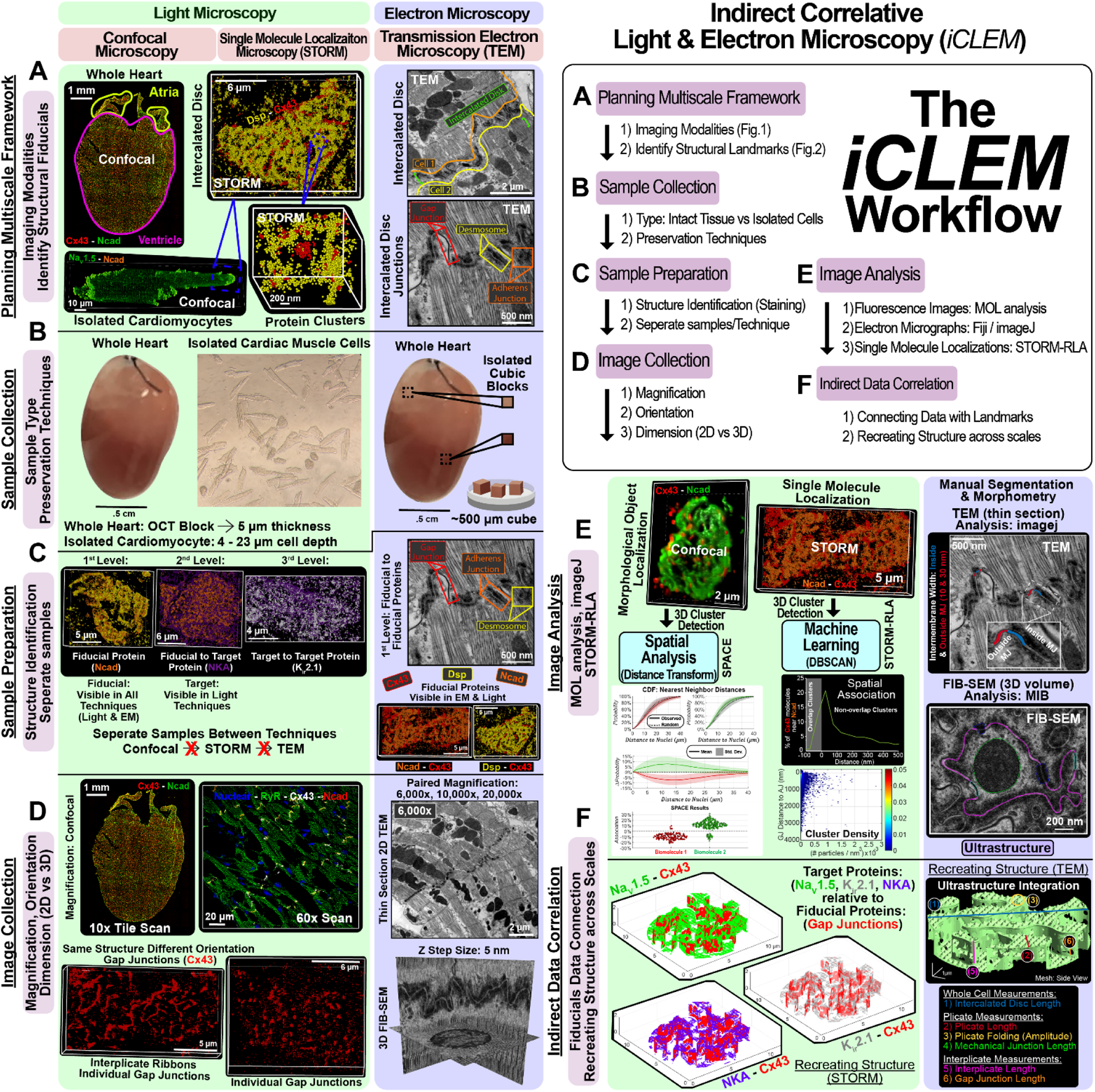
Indirect Correlative Light & Electron Microscopy (*iCLEM*) Workflow. Simplified flowchart (top right) and example images illustrating the steps involved in *iCLEM*: **A)** Planning, **B)** Sample collection, **C)** Sample preparation, **D)** Image collection, **E)** Image analysis and **F)** Indirect correlation of morphometric measurements.

**Figure 2.**
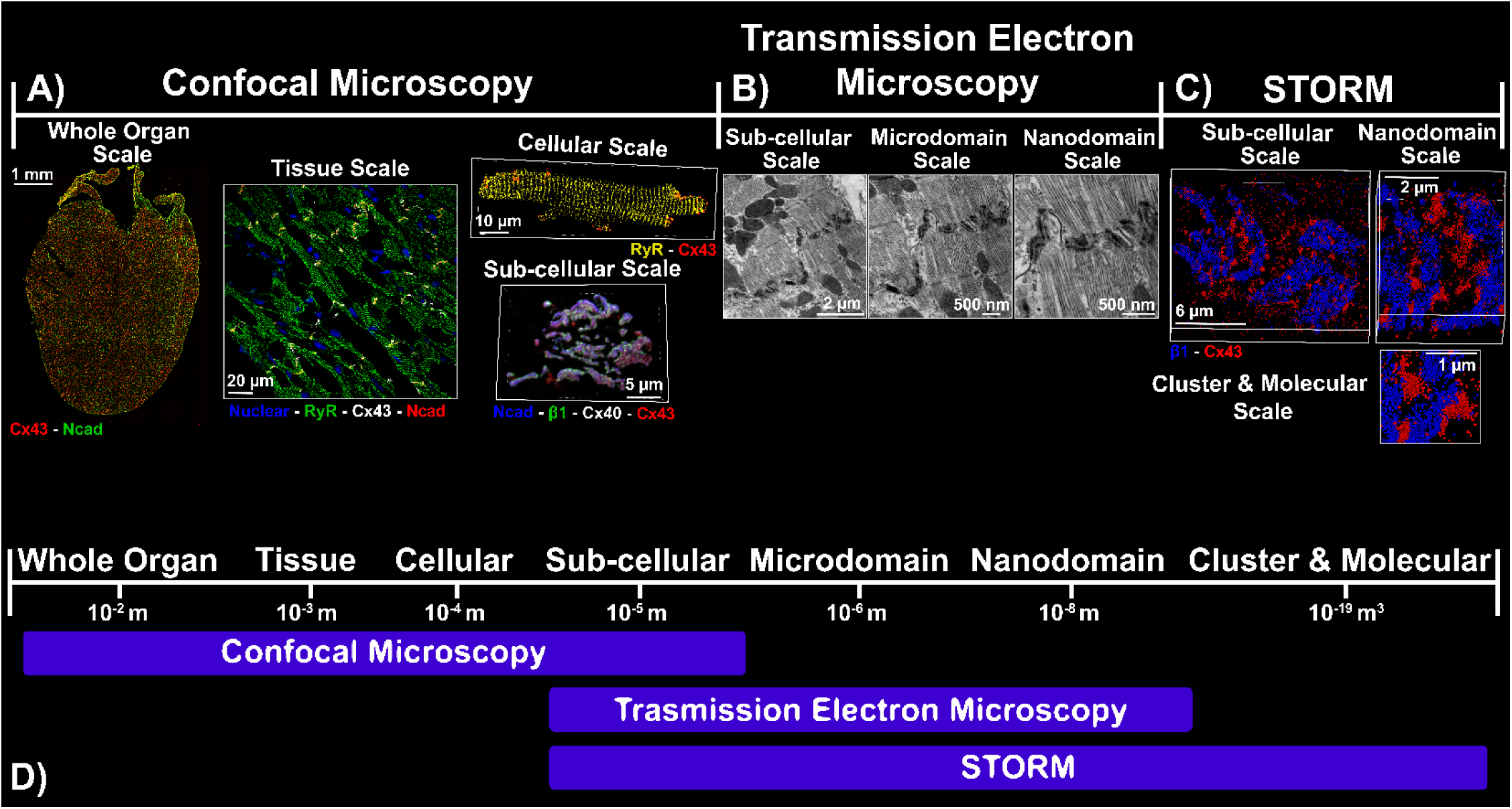
*iCLEM* Microscopy Techniques. Schematic illustrating some of the specific imaging modalities incorporated into the implementation of *iCLEM* described here indicating the spatial scales accessible to each.

### Planning Multiscale Experimental Framework

In the example shown in Figure 2, cardiac muscle tissue was imaged using light and electron microscopy techniques in order to characterize the heterogeneous structural properties of specialized cell-cell contact sites between cardiac muscle cells, known as intercalated discs. Specific imaging modalities were selected based on their ability to resolve and capture structures ranging in scale from the whole organ to cluster and molecular levels. Confocal microscopy enables visualization of fluorescently labeled biomolecules within the intercalated disc, albeit with diffraction-limited resolution, covering the whole organ (heart), tissue, cellular (ventricular cardiac muscle cell), and sub-cellular scales (intercalated disc; Figure 2A). TEM provides exquisite ultrastructural detail, covering sub-cellular (intercalated disc), microdomain, and nanodomain scales (Figure 2B, 5B). STORM enables studies below the diffraction limit (attaining ∼20 nm lateral and <50 nm axial resolution), and covers sub-cellular (intercalated disc), nanodomain, and cluster/molecule scales (Figure 2C, 7B). Importantly, experiments were designed to enable cross-validation of structural findings at multiple levels by exploiting degrees of overlap in scales / structures captured by different imaging modalities (Figure 2D). Samples were prepared and imaged separately for each imaging modality employed.

### Selection and Use of Intrinsic Fiducials

At the heart of *iCLEM* are intrinsic fiducials – structures identifiable between light and electron microscopy – which served as reference structures relative which morphometric measurements could be made to enable correlation of data across imaging modalities (Figure 3). In addition to being apparent via different imaging modalities, intrinsic fiducials are also selected with some consideration towards their ability to demarcate specific structural regions at larger spatial scales, where the fiducial itself may not be fully resolvable (Figure 3A-H). Thus, in this example, three types of cell-cell junctions located within intercalated discs were selected as intrinsic fiducials (Figure 3E, F, I): 1) gap junctions, composed of connexin 43 (Cx43, red), 2) adherens junctions, composed of N-cadherin (N-cad, orange), and 3) desmosomes, containing desmoplakin (Dsp, yellow)(Nielsen, et al., 2023; Vermij, et al., 2017). Stepping up to larger scales, adherens junctions and desmosomes predominantly occupy highly folded intercalated disc membranes oriented perpendicular to the cell’s long axis, known as plicate microdomains (Figure 3H, purple), while gap junctions are predominantly localized to intervening interplicate microdomains (Figure 3H, cyan), which run parallel to the cell’s long axis(Nielsen, et al., 2023; Severs, 1990; Struckman, et al., 2023). Thus, at these scales N-cad and Dsp serve as molecular landmarks for plicate microdomains, while Cx43 marks interplicate microdomains (Figure 3D). At cellular through organ scales (Figure 3A-C), different intercalated disc subdomains cannot be clearly discriminated; however, the cell-cell junctions, and thereby, the corresponding molecular fiducials (Cx43, N-cad, Dsp) serve as landmarks for the longitudinal ends of cardiac muscle cells, the predominant location of intercalated discs(Agullo-Pascual, et al., 2014; Makara, et al., 2014; Mohler, et al., 2004). Additionally, contextual anatomical cues gleaned from visualizing intercalated discs can help discriminate between larger scale feature such as the upper (atria) and lower (ventricles) chambers of the heart (Figure 3A).

**Figure 3.**
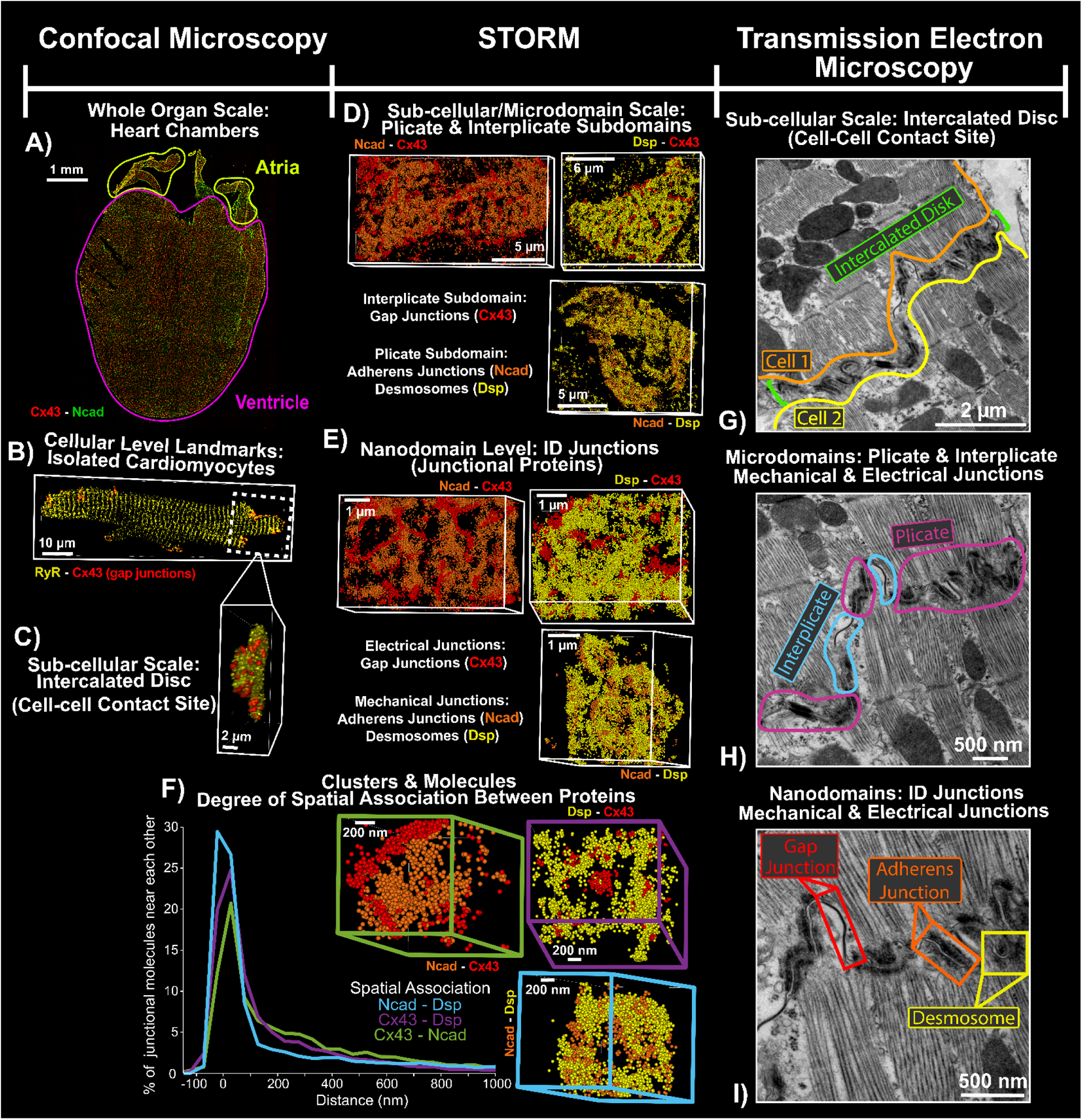
Intrinsic Structural Fiducials. *iCLEM* exploits intrinsic fiducials – structures identifiable between light and electron microscopy – to correlate independently collected morphometric measurements from individual imaging modalities. In the example shown here, gap junctions/Cx43 protein, adherens junctions/N-cad protein and desmosomes/Dsp protein are exploited as intrinsic fiducials to capture positional context at different spatial scales. **A-C)** Use of confocal microscopy at different scales. Range of spatial scales at which features are captured by **D-F)** single molecule localization microscopy and **G-I)** transmission electron microscopy.

### Sample Handling, Imaging, and Analysis

#### Confocal Microscopy

Samples for confocal microscopy consisted of 5 µm thick cryosections obtained from fresh-frozen cardiac tissue and freshly isolated primary cardiac muscle cells adhered on to glass coverslips, which were fixed using paraformaldehyde (Figure 1B-C, Figure 4). A subset of samples was immunolabeled to detect fiducial biomolecules (Cx43, N-cad, Dsp) relative to each other to enable cross-validation with TEM-derived ultrastructural measurements, while the remainder were immunolabeled to detect one or more target biomolecules (not identifiable ultrastructurally without immunolabeling) in relation to one or more fiducial biomolecules (Figure 4). Image collection utilized different workflows, each optimized to capture features at specific spatial scales. Tilescan imaging using a 10x/0.45NA objective was used to image entire frontal sections of mouse hearts (Figure 4B, left). Additionally, close-up views of smaller regions of tissues (< 200 x 200 µm) and individual intercalated discs (Figure 4B, right) were obtained as single plane images (for survey views only) or z-stacks (images used for quantitative analysis) using a 60x/1.4NA objective. These data enabled structural studies at scales ranging from whole heart (differences between upper and lower chambers) down to the level of intercalated disc microdomains (Figure 4A), which were the smallest features identifiable using diffraction-limited confocal microscopy (lateral resolution 200-300 nm). Additionally, to circumvent challenges associated with effectively/efficiently segmenting individual muscle cells within tissue images, confocal imaging was performed on cardiac muscle cells isolated from the upper (atria) and lower chambers (ventricles) to assess the distribution of target biomolecules (in this example: the cardiac isoform of the voltage-gated sodium channel, Na_V_1.5) between the intercalated disc and other membrane sites in cardiac muscle cells using the adherens junction protein N-cad as a marker for intercalated discs (Figure 4C). Individual cells were imaged as z-stacks using a 60x/1.4NA objective. For all confocal microscopy studies, fluorophores were sequentially excited and imaged to minimize cross-channel bleed-through effects.

**Figure 4.**
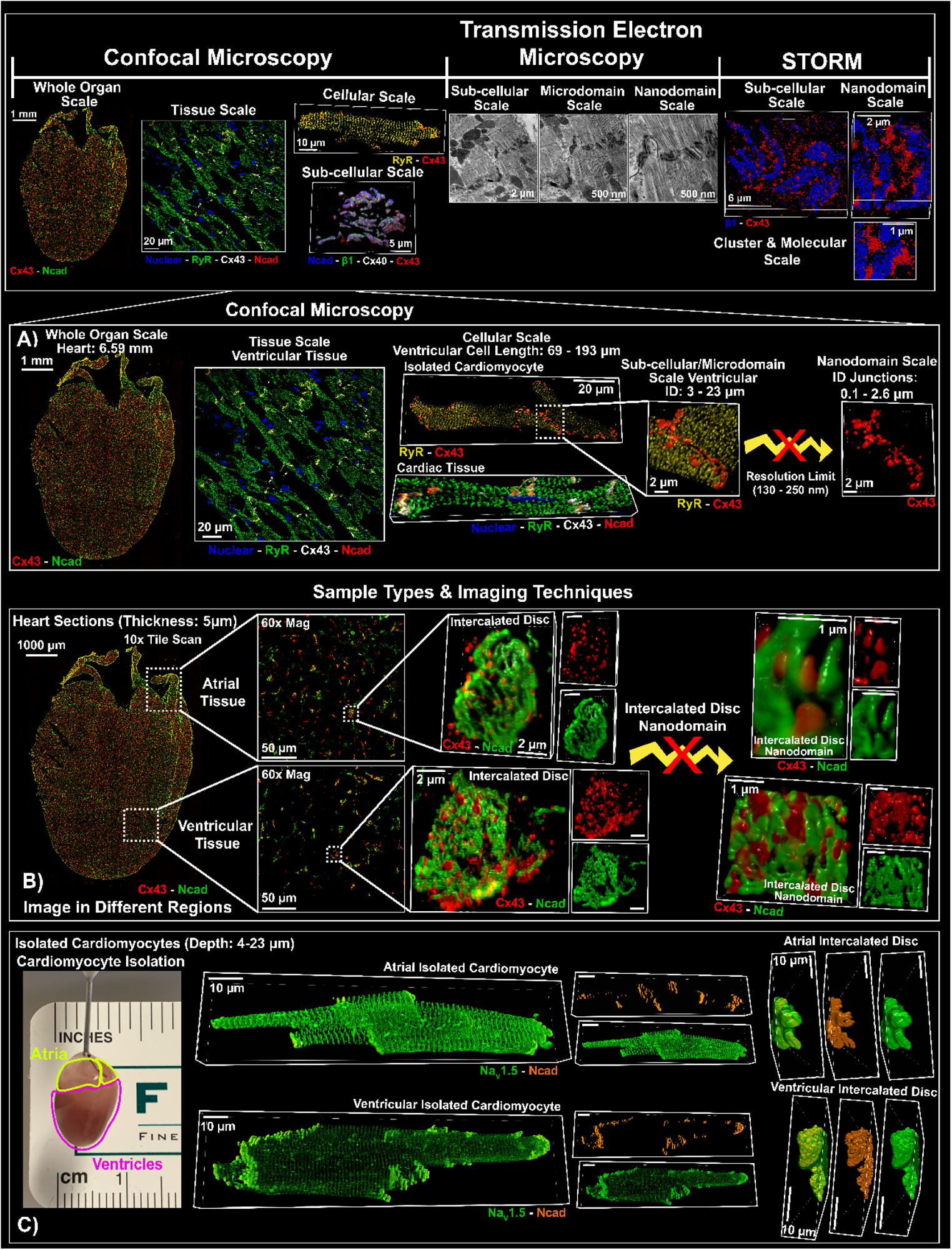
Confocal Microscopy. Overview of imaging modalities incorporated into the *iCLEM* implementation in the present study (top) and the various uses of confocal microscopy therein, illustrating **A)** spatial scales covered, and implementation with **B)** cryosections of heart tissue and **C)** isolated cardiac muscle cells.

As noted above, confocal images were analyzed using our previously published point process-based spatial analysis pipeline, Spatial Pattern Analysis using Closest Events (SPACE; Figure 1E)(Bogdanov, et al., 2021). This approach, which can be applied to any digital image, regardless of imaging modality, uses distance transformations to efficiently measure the nearest neighbor distances for segmented immunosignals for a given biomolecule in relation to sub-cellular landmarks (eg. nuclei, cell periphery) or a second, co-labeled biomolecule. By comparing the distribution of observed nearest neighbor distances with that predicted under complete spatial randomness [an approach similar to statistical F and G functions(Unwin, 1996; Ying, 2013)], quantitative measurements are obtained for the strength of non-random attraction / repulsion between co-labeled molecules / structures. Importantly, nearest neighbor distance distributions for a target biomolecule in relation to fiducial biomolecule allow the spatial distribution of the target biomolecule to be placed in the context of ultrastructural properties at such locations (morphometric measurements at sites specified in relation to fiducial structures in electron micrographs obtained from separate samples).

#### Electron microscopy

Tissue samples for electron microscopy were collected as tissue blocks (∼500 μm) fixed in 2.5% glutaraldehyde (Figure 1B-C). Tissue blocks were processed, sectioned (70 nm) and stained for TEM imaging (Figure 5C). Paired sets of images were collected of each identified intercalated disc at multiple magnifications to capture features at sub-cellular, microdomain, and nanodomain scales. These 2D data were supplemented by 3D data obtained using volume electron microscopy (specifically, FIB-SEM) performed at 10,000x magnification with 5 nm z steps between sequential images on tissue blocks processed and stained as described above (Figure 5D, 6B).

**Figure 5.**
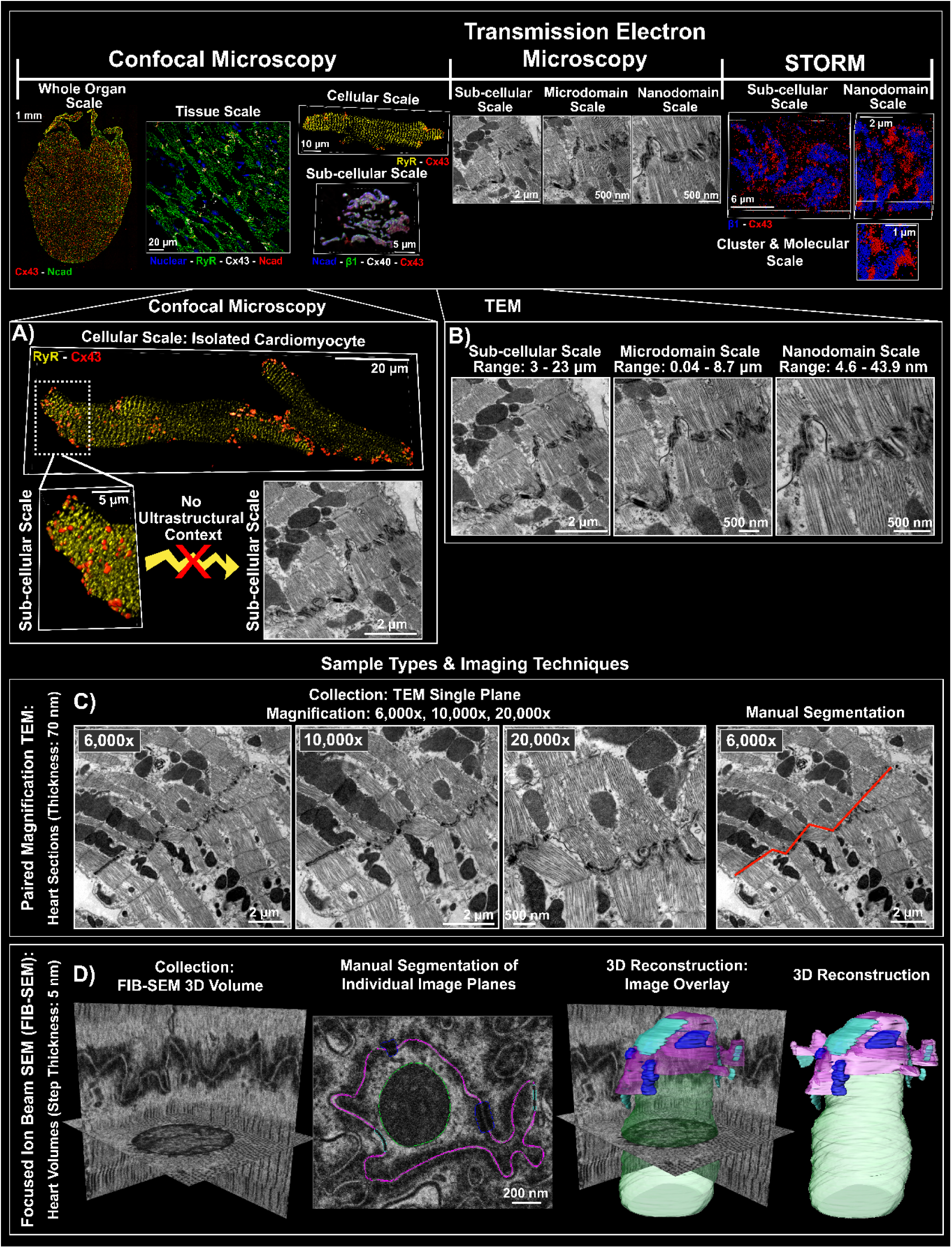
Electron Microscopy. Overview of imaging modalities incorporated into the *iCLEM* implementation in the present study (top) and the various uses of electron microscopy therein, illustrating **A, B)** spatial scales covered by electron microscopy in comparison with confocal microscopy, and implementation with **C)** TEM for 2D imaging and **D)** FIB-SEM for volume EM.

Although progress is being made on automating segmentation of electron micrographs, these methods remain limited to specific use cases and the resulting segmentation pipelines lack versatility. Therefore, we opted to use manual segmentation and morphometry of TEM images performed using ImageJ as previously described (Figure 1E)(Moise, et al., 2021; Struckman, et al., 2023). Paired analysis of multiple images of the same structure (in this case, intercalated discs) obtained at different magnifications enabled quantification and correlation of ultrastructural properties at different spatial scales (Figure 6A): 1) Sub-cellular scale: cell width at the intercalated disc and total intercalated disc length (cross-sectional length); 2) Microdomain scale: dimensions of intercalated disc subdomains, plicate and interplicate, amplitude and frequency of plicate membrane waviness, lengths of gap junctions (GJ) and mechanical junctions (MJ); and 3) Nanodomain scale: intermembrane distances inside adherens junctions / desmosomes and at juxta-junctional sites. Together, such measurements provided seamless quantification of ultrastructure from micro-through nano-scales, enabling the generation of realistic finite element models (Figure 1F).

**Figure 6.**
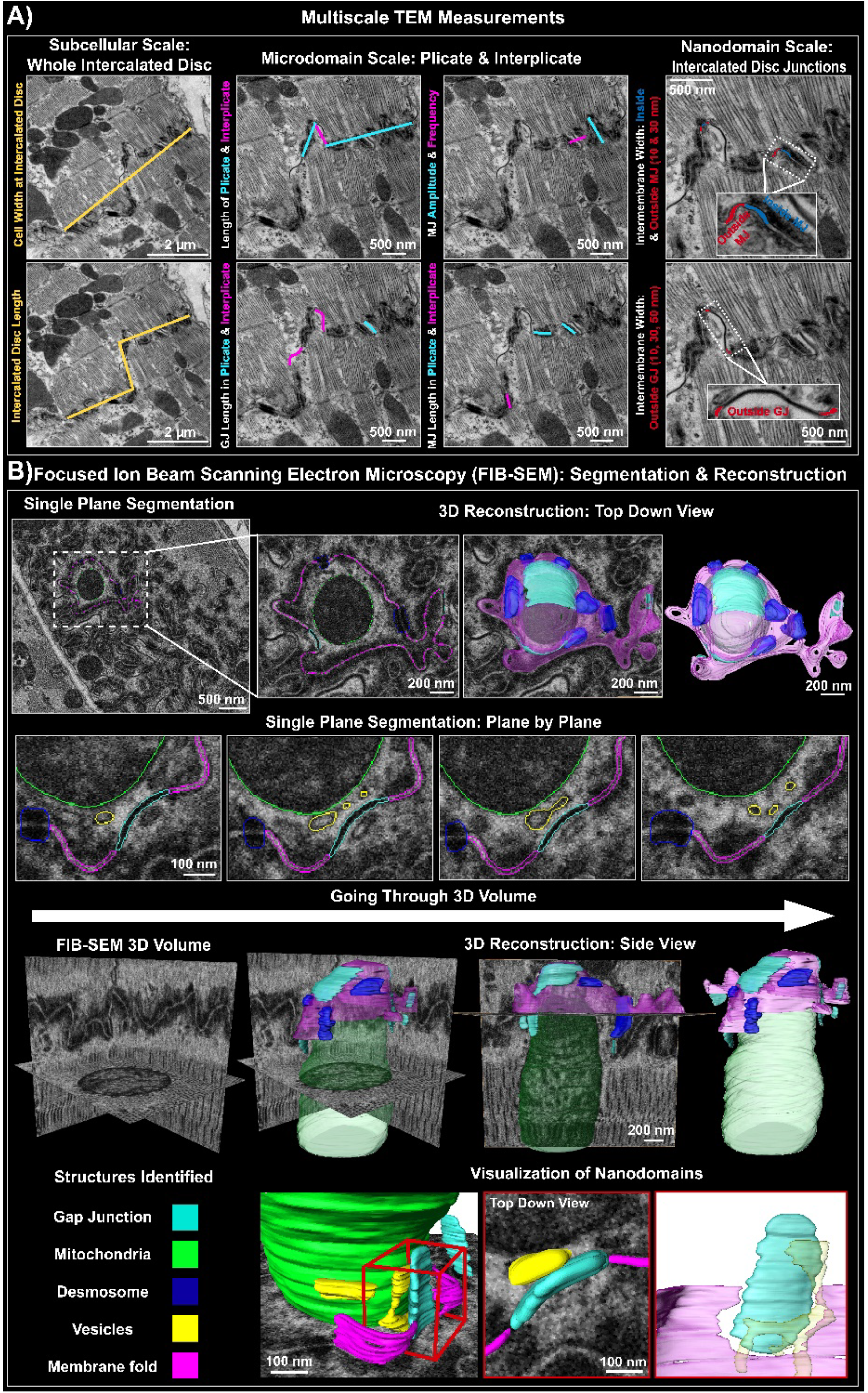
*iCLEM* Application Example: Electron Microscopy. Specific uses of electron microscopy in the *iCLEM* presented herein illustrating **A)** multiscale ultrastructural quantification from paired sets of TEM images at different magnifications and **B)** use of FIB-SEM to capture 3D ultrastructural features. Representative 2D and 3D FIB-SEM images show computational reconstruction of segmented structures overlaid on raw image data.

Manual plane-by-plane segmentation of multiplane 3D volume image stacks from FIB-SEM was performed using Microscopy Image Browser (Figure 1E, 5D, 6B) as previously described(Vanslembrouck, et al., 2018). Segmentation was performed on images collected at 10,000x magnification to capture nanodomain level structural features (Figure 6B). Color-coded annotations of segmented structures were overlaid on single plane images (top down view) and 3D volume stacks (side view) to verify faithful reconstruction (Figure 6B).

#### Single Molecule Localization Microscopy

Cell and tissue samples for single molecule localization microscopy were collected and processed as for confocal microscopy with specific modifications to the protocol as noted above (Figures 1B-C, 7). Cells and tissue sections were adhered on to glass coverslips and STORM imaging was performed in widefield configuration as described above. Given our ability to only visualize two biomolecules within a given sample, images were collected of 1) intrinsic fiducial biomolecules relative to each other (enabling cross-validation with TEM-derived ultrastructural measurements), 2) target biomolecules relative to fiducial biomolecules (thus placing target biomolecules within ultrastructural context) and 3) target biomolecules relative to each other. In the example shown in Figure 7C, 1) the fiducial protein N-cad (indicative of adherens junctions) was first localized in relation to the fiducial protein Dsp (indicative of desmosomes), 2) then the target protein NKA (sodium/potassium ATPase) was localized in relation to the fiducial protein N-cad, and 3) finally, the target protein K_ir_2.1 (inward-rectifier potassium channel) was localized relative to NKA. It should be noted here that these three steps were accomplished using independent sets of samples and distributions of the resulting measurements correlated in keeping with the operating principles of *iCLEM*. Figure 8 illustrates a full implementation of this approach, assessing the localization of 1) fiducial proteins (Cx43, N-cad and Dsp) relative to each other, 2) target proteins, Na_V_1.5 (cardiac isoform of the voltage-gated sodium channel), K_ir_2.1 (inward rectifying potassium channel), and NKA (sodium-potassium ATPase) relative to the fiducial proteins, and finally, 3) the target proteins relative to each other to obtain a comprehensive understanding of intercalated disc molecular organization.

**Figure 7.**
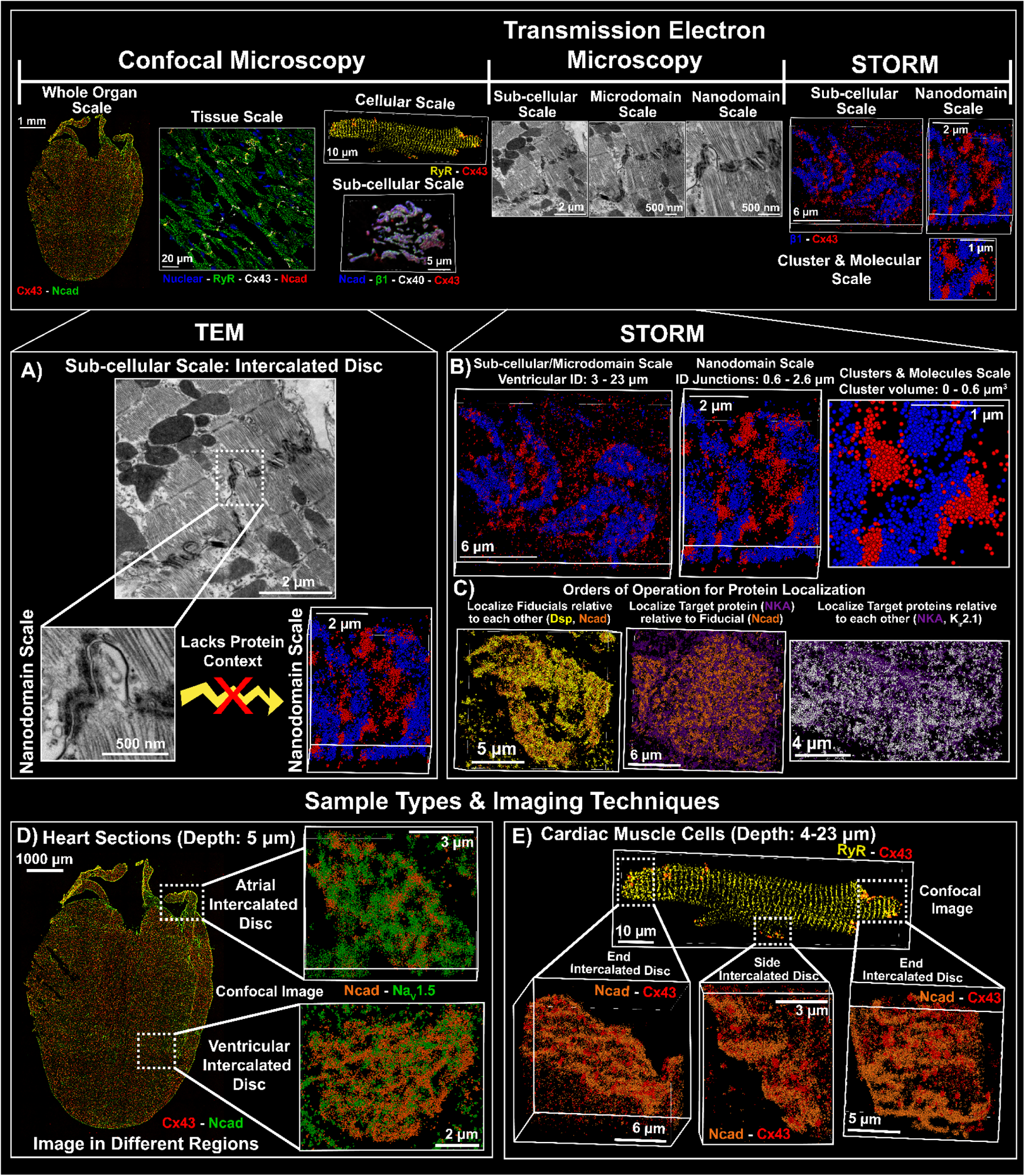
STochastic Optical Reconstruction Microscopy (STORM). Overview of imaging modalities incorporated into the *iCLEM* implementation in the present study (top) and the various uses of single molecule localization microscopy therein, illustrating **A, B)** spatial scales covered by electron microscopy in comparison with STORM. **C)** Use of intrinsic fiducials to systematically localize target proteins. STORM imaging of intercalated discs in **D)** tissue sections and **E)** isolated cardiac muscle cells.

**Figure 8.**
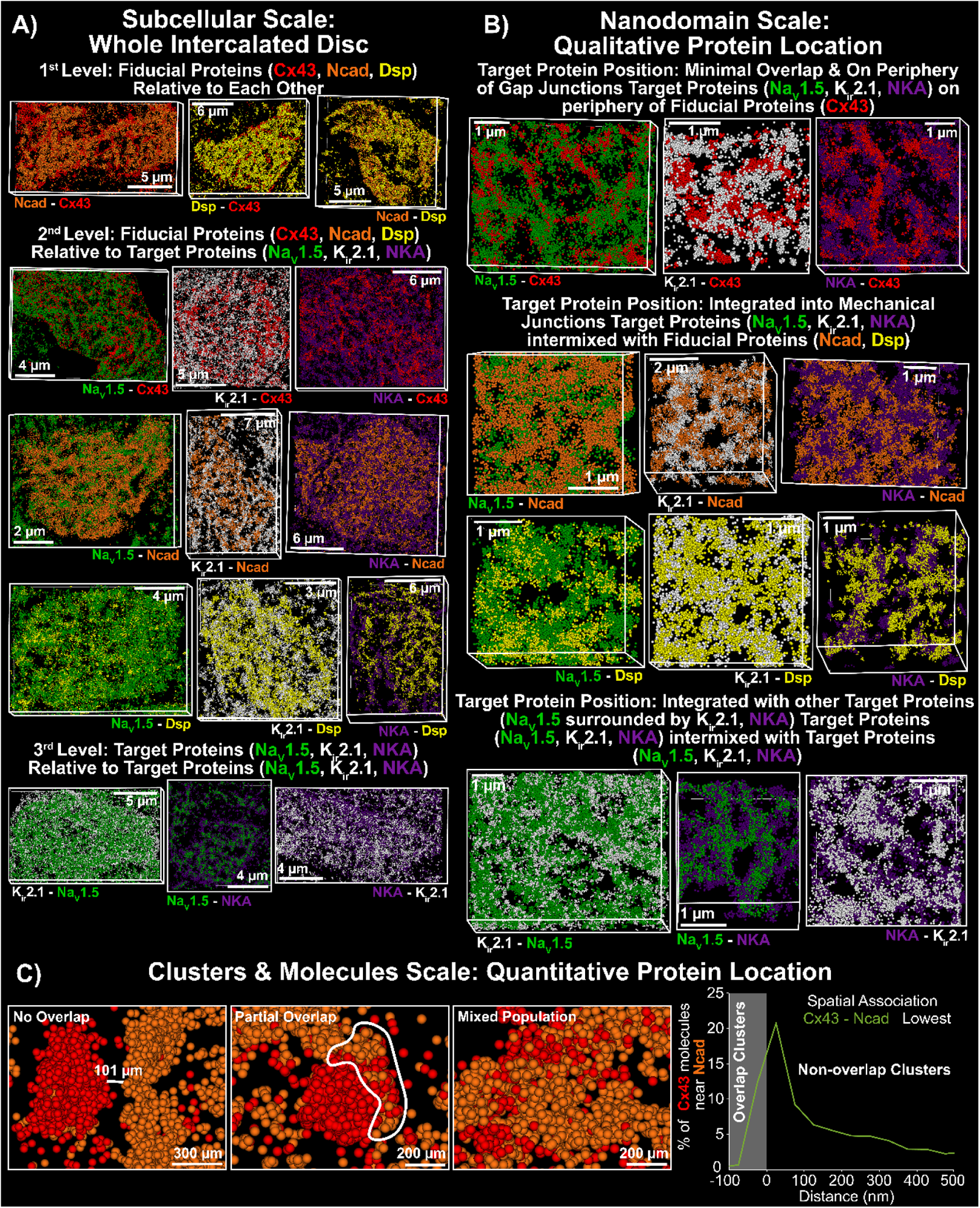
*iCLEM* Application Example: STORM. Representative STORM images showing **A)** whole intercalated discs and **B)** zoomed in views of individual clusters illustrating the technique’s incorporation into the *iCLEM* pipeline progressing from localizing fiducial proteins relative to each other to target proteins relative to fiducial proteins and finally target proteins relative to each other. **C)** Representative images illustrating different levels of inter-cluster association and a sample probability density function of inter-cluster nearest neighbor distances between gap junctions (Cx43) and adherens junctions (Ncad).

In tissue sections, intercalated discs could be observed in a variety of orientations ranging from edgewise (oriented along an axial plane) to *en face* (oriented in the lateral plane) (Figure 1D, 7D). In order to maximize the ability to resolve distinct subdomains within the intercalated discs, all images from tissue samples were of intercalated discs in *en face* orientation. In contrast, isolated cardiac muscle cells were oriented with their long axis parallel to the imaging plane by virtue of their position on the coverslips, requiring intercalated discs to be reconstructed along an axial (xz / yz) plane (Figure 7E). Of note, multi-focus imaging rather than astigmatism was used for localizing the axial position of fluorophores due to the former’s superior performance in thicker, more optically dense samples(Juette, et al., 2008; Liu, et al., 2020).

Single molecule localizations were analyzed using our previously published STORM-based Relative Localization Analysis (STORM-RLA(Veeraraghavan & Gourdie, 2016)), a machine learning-based cluster analysis approach that uses density-based spatial clustering for applications with noise (DBSCAN) as the underlying clustering algorithm. By fitting surfaces to detected clusters, this approach yields distributions of nearest neighbor distances between clusters of co-labeled biomolecules (Figure 8C, right), which can then be subjected to spatial statistical analysis and correlated with electron microscopy-derived ultrastructural measurements as detailed above. It should be noted here that this analysis can be applied to any set of localizations in continuous space, regardless of the specific imaging modality used.

#### Indirect Correlation of Measurements and Integration with Computational Modeling

Indirect correlation of morphometric measurements from electron microscopy and protein localization from light microscopy is achieved by relating the distributions of measures using their spatial positions recorded in relation to intrinsic fiducials. For example, distributions of intermembrane distances (from electron microscopy; Figure 6A), measured at various positions relative to the edges of gap junctions, desmosomes, and adherens junctions, are correlated with various properties (mass, volume, density; Figure 8C) of target protein clusters and their spatial association with each other at such sites (from light microscopy). To more, directly probe the functional implications of observed structural properties, we generated populations of finite element models of intercalated discs based on electron microscopy-derived ultrastructure (Figure 9, top), following our recently published approach(Moise, et al., 2021). Next, these meshes were populated with target proteins, which were placed into the meshes in relation to intrinsic fiducials (gap junctions, adherens junctions, desmosomes)(Struckman, et al., 2023). The completed intercalated disc meshes were then incorporated into physiological models of electrical excitation spread through chains of cardiac muscle cells to generate predictions of impulse conduction velocity and other functional descriptors of cell-cell communication (Figure 9, bottom). These predictions were then validated against experimental measures obtained from whole-heart functional imaging studies conducted by labeling tissues with a voltage-sensitive dye to enable direct observation of transmembrane potentials (Figure 9, bottom right), as previously described(King, et al., 2021). In a recent study, we applied this approach to uncover previously unanticipated structure-function relationships underpinning cell-cell communication in the heart(Struckman, et al., 2023), helping identify novel targets for mechanism-based therapy to prevent cardiac rhythm disturbances resulting from impaired cell-cell communication.

**Figure 9.**
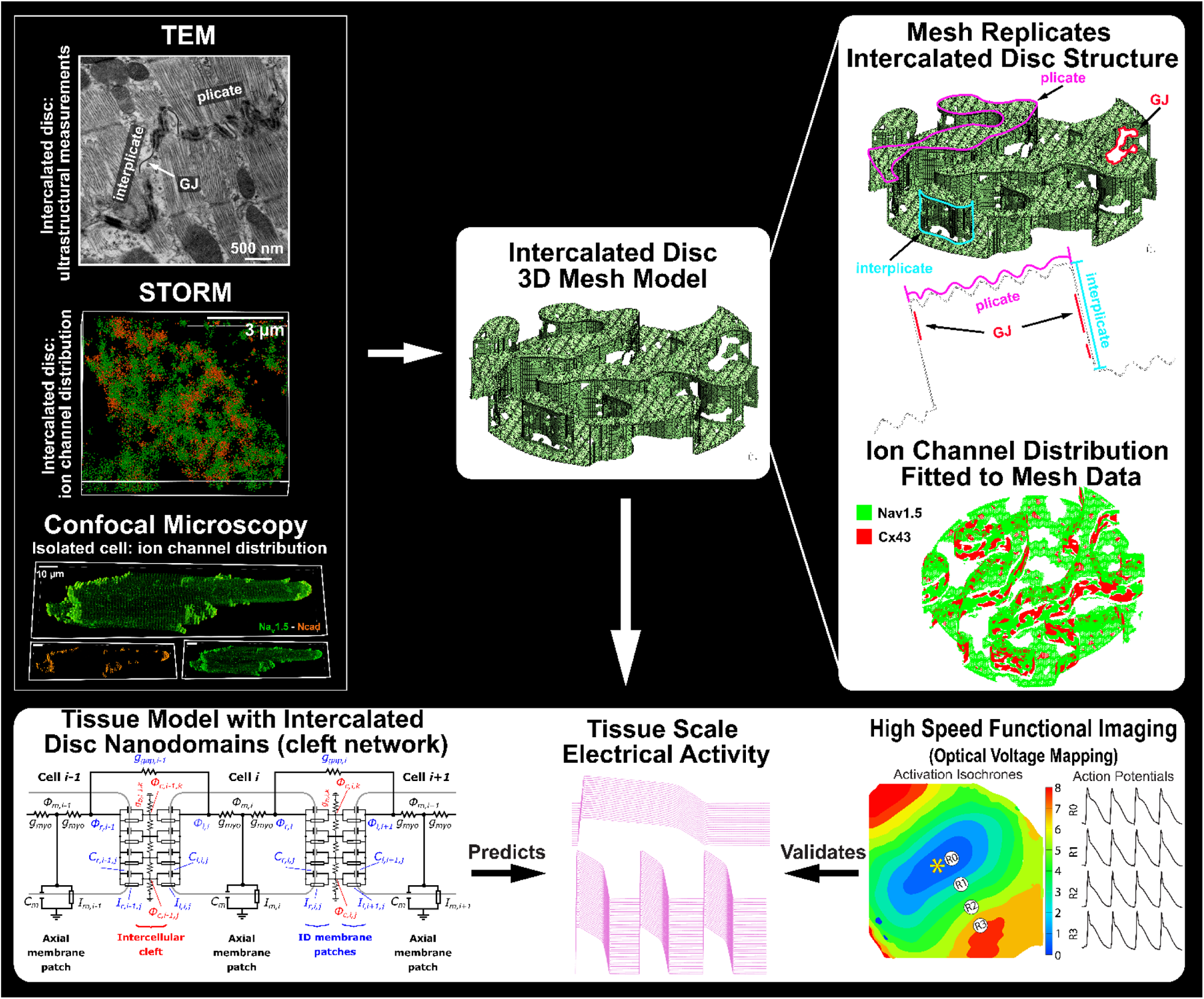
Integration with Computational Modeling. Schematic showing integration of light microscopy-derived protein localization into finite element models of intercalated discs constructed using electron microscopy-derived ultrastructural measurements (top) and the use of such models to predict function properties (bottom), which can be validated against macroscopic experimental data (bottom).

## Discussion

The study of structure remains one of the most valuable approaches to understanding the function of complex biological systems. Thus, structural microscopy continues to grow in value in the life sciences. Historically, electron microscopy offered life scientists exquisite spatial resolution and rich ultrastructural context, albeit with very limited ability to identify specific biomolecules within that ultrastructural context. In contrast, fluorescent light microscopy renders it incredibly easy to label and visualize specific biomolecules, albeit with minimal, if any, ultrastructural context, and with spatial resolution limited to 200-300 nm by diffraction. Although super-resolution light microscopy methods have improved spatial resolution by an order of magnitude (Jacquemet, et al., 2020), the lack of ultrastructural context remains a barrier to be overcome. Additionally, with both light and electron microscopy techniques, high spatial resolution often comes at the cost of a restricted field of view, limiting the ability to assess structural properties across a wide range of spatial scales. As a result, available data on the structure of cells and tissues is often fragmentary and restricted to specific spatial scales, limiting the biologist’s ability to collate knowledge across scales.

Against this backdrop, correlative light and electron microscopy (*CLEM*) stands poised to revolutionize the life sciences by placing molecular organization gleaned from light microscopy in the context of ultrastructure visualized using electron microscopy(Friedrichsen, et al., 2022; Hegermann, et al., 2019; Oorschot, et al., 2021; Perkovic, et al., 2014). However, this entails preparing samples to meet the disparate chemistry criteria imposed by light and electron microscopes and then obtaining precisely registered images of nanoscale neighborhoods within a sample on multiple microscopes(Ferguson, et al., 2017; Kremer, et al., 2021; Yang, et al., 2021). Thus, direct CLEM approaches require extremely high cost, skill and time commitments, and have far lower (10-100 fold) throughput (images generated per person-hour of effort) than either light or electron microscopy alone. These factors not only severely restrict the adoption of CLEM methods but also preclude their use on precious samples and to quantify rare / subtle effects (by assessing intra- and inter-sample heterogeneities), such as remodeling associated with early stages of disease. Additionally, the field-of-view of any direct CLEM techniques is limited to the smallest afforded by the specific light and electron microscopy modalities used, limiting their applicability to multiscale investigations.

We developed *indirect* CLEM (*iCLEM*) to complement direct CLEM methods and specifically address the aforementioned challenges. Unlike direct CLEM approaches, *iCLEM* entails the preparation and imaging of separate samples on two or more light and electron microscopes followed by indirect correlation of morphometric measurements vis-à-vis intrinsic fiducials identifiable via all the imaging modalities employed. Thus, *iCLEM* is agnostic to the specific imaging modalities utilized and can be customized based on the demands of the investigation and the resources available to the investigators. It should be noted here that direct CLEM methods use extrinsic fiducials (beads or grids) added to the sample to enable registration of images across imaging modalities, whereas *iCLEM* uses intrinsic fiducials as reference points for measurements made from disparate sets of samples prepared for each imaging modality, with correlation accomplished between populations of structures studied rather than in a one-to-one fashion. Provided appropriate intrinsic fiducials can be identified, *iCLEM* preserves the advantages inherent to each imaging modality employed in terms of ease, throughput and cost effectiveness of sample preparation and image collection without the need for highly specialized (and thus, not widely available) hardware tools. Additionally, *iCLEM* could also serve as a low cost option for investigators precluded by resource limitations from using direct CLEM methods as well as helping to identify specific, high priority experiments that merit the resource commitment demanded by direct CLEM studies. Thus, *iCLEM* serves as a complement, rather than an alternative, to direct CLEM approaches, extending the biologist’s reach to capture subtle phenomena across a wide range of spatial scales.

As with any experimental modality, the benefits afforded by *iCLEM* come at a cost. In addition to the inability to directly correlate image-to-image, *iCLEM* requires prior knowledge of the cells/tissues being studied for appropriate intrinsic fiducials to be identified. Given that these intrinsic fiducials are used to correlate morphometric measurements across imaging modalities, they cannot be substituted by extrinsic fiducials (grids, beads), which are only useful for image registration when the same sample is imaged using different modalities. *iCLEM* also demands the collection of datasets large enough to fully characterize the distributions of measured structural properties and their intra- and inter-sample heterogeneities. In the examples provided above, we demonstrate the *iCLEM* approach, combining two light microscopy techniques (confocal and STORM) with two electron microcopy methods (TEM, FIB-SEM) to characterize the structure of cell-cell contacts in the heart, termed intercalated discs, across a wide range of spatial scales. Whereas we are unable to correlate molecular organization from STORM with ultrastructure from TEM for any specific intercalated disc, we are able to assess heterogeneities in structural properties and correlate them across populations of intercalated discs. In other words, direct image-to-image correlation provides the fullest understanding of a specific structure observed, while indirect correlation enables unprecedented quantification of structural properties across the population of structures observed. We further demonstrate how *iCLEM*-derived quantitative data can be readily integrated into computational models to uncover structure-function relationships underlying the behaviors of biological systems, further extending the investigative power of structural microscopy. This is demonstrated in a recent study where we utilized *iCLEM* to unravel chamber-specific differences in intercalated disc ultrastructure and molecular organization and their impact on the spread of electrical signals through the heart(Struckman, et al., 2023).

## Summary

In summary, we demonstrate here an indirect CLEM approach, which exploits structural fiducials inherent within samples to circumvent the need for the same sample to be imaged using multiple modalities. This approach represents an important step towards democratizing CLEM, lowering the cost, expertise, and time commitments demanded by direct CLEM techniques, while enabling unprecedented quantitative assessment of structural properties and their heterogeneities across a wide range of spatial scales. Additionally, *iCLEM* is agnostic to the specific imaging modalities employed, enabling investigators to customize implementation to meet the demands of specific investigations.

## Acknowledgements

The authors would like to thank Ms. Sarah Mikula, and Mr. Jeff Tonniges from the OSU Campus Microscopy & Imaging Facility for preparing the electron microscopy samples for this study.

## Declarations

### Availability of data and material

Raw experimental data are backed up on OSU servers and will be shared freely on request.

### Author Contributions

H. L. Struckman: performed light microscopy and transmission electron microscopy experiments, helped develop the *iCLEM* concept, and prepared the manuscript.

N. Moise: performed computational modeling, and helped with manuscript preparation.

B. Vanslembrouck: performed volume electron microscopy experiments and assisted with manuscript preparation.

N. Rothacker: assisted with light microscopy studies.

Z. Chen: generated and validated custom anitbodies used in this study and assisted with interpretation of results.

J. van Hengel: oversaw the volume electron microscopy studies and assisted with interpretation of results and manuscript preparation.

S. H. Weinberg: oversaw the computational modeling studies and assisted with interpretation of results and manuscript preparation.

R. Veeraraghavan: conceived of the *iCLEM* concept oversaw the research project, developed and implemented image analysis, and assisted with manuscript preparation. Corresponding author.

### Ethics approvals

All animal procedures were approved by Institutional Animal Care and Use Committee at The Ohio State University and performed in accordance with the Guide for the Care and Use of Laboratory Animals published by the U.S. National Institutes of Health (NIH Publication No. 85-23, revised 2011).

